# Fish remain high in selenium long after mountaintop coal mines close

**DOI:** 10.1101/2025.05.22.655156

**Authors:** Colin A. Cooke, Jennifer A. Graydon, Andreas Luek, Xiufen Lu, Hailey Yu, X. Chris Le, Megan Reichert

## Abstract

Mountaintop removal (MTR) coal mining generates large volumes of waste rock. Weathering of this waste rock releases selenium, which can bioaccumulate to levels that can harm, and even extirpate, downstream fish communities. This is well demonstrated in ecosystems impacted by active MTR operations; however, less is known about the long-term impacts after coal mines close. Here we show that MTR coal mines still present an acute threat to downstream fish populations, decades after mining ends. Crowsnest Lake (Alberta, Canada) receives runoff from the Tent Mountain Coal Mine, which closed in the 1980s and part of which was certified reclaimed. Fish in Crowsnest Lake contain tissue selenium concentrations (5–26 µg/g dry weight) that exceed guidelines and rival fish selenium levels downstream of active MTR operations. This is despite lake water selenium concentrations (≤2 µg/L) that are below water quality guidelines intended to protect fish. The clinical signs of selenium poisoning in fish are similar to the symptoms of Whirling Disease, which was first detected in the Crowsnest basin in 2016 making this the first aquatic system to be impacted by both stressors. This finding demonstrates that the biological impacts of MTR coal mining can persist long after mining operations end, and it suggests that any further coal mine development may well push the Crowsnest fishery beyond sustainability.

**Synopsis:** Fish in Crowsnest Lake, Alberta, are high in selenium sourced from an abandoned mountaintop removal coal mine.

## Introduction

Coal mining in the Canadian Rocky Mountains has evolved from underground to modern mountaintop removal (MTR) methods. MTR is a type of surface mining at the summit (or along a summit ridge) of a mountain that removes overburden to expose coal deposits.^1–3^ MTR creates a waste rock problem, and leaching of material from waste rock piles can pollute downstream ecosystems. Previous monitoring and research have identified elevated concentrations of selenium^1,4–8^, nitrates^1,6,9^, and ions^2,6–8^ in rivers draining mountaintop removal coal mines. The inputs of these pollutants can have devastating consequences for downstream fish communities and the ecosystems they inhabit. This is especially true for selenium, which bioaccumulates and biomagnifies through foodwebs. Selenium is an essential micronutrient that at higher concentrations has toxic effects. In fish, teratogenic effects lead to embryonic malformations when selenium is transferred into the eggs during ovarian development. This results in reduced reproductive success up to complete reproductive failure in exposed fish populations. While the effects on newly hatched fish may be rapid and severe (i.e., acute), the parent fish may appear unaffected. Complete reproductive failure and even population collapse can occur if exposure is prolonged and high enough.

The Crowsnest Pass spans the Continental Divide separating Pacific from Atlantic drainage basins (and British Columbia from Alberta) in western Canada. On the Alberta side of the pass, underground coal mining began in late 1800s, and the first MTR mine (Tent Mountain) opened in 1948. A second MTR mine (Grassy Mountain) opened in 1956; by 1983 both mines had (and remain) closed. A 2016 application to mine the remaining coal generated significant debate.^15^ The application was eventually rejected by a joint Provincial-Federal review panel in 2021, which cited significant adverse environmental effects that outweighed any positive economic impacts of the project.^19^ The potential for elevated selenium pollution was paramount in this decision, and any new selenium inputs would be additive to existing inputs from the legacy coal mines. These legacy inputs were quantified in a recent study that revealed high selenium concentrations in the streams and creeks draining the abandoned Tent and Grassy Mountain MTR mines^8^. Here, we present a first assessment of metals in fish from Crowsnest Lake. Our new results reveal these fish already contain high selenium concentrations that rival populations impacted by active MTR mining.

## Methods

### Study region and regional water quality

The Crowsnest River valley contains multiple towns and it is a major transportation corridor, with Highway 3 and the Canadian Pacific Railway running parallel to Crowsnest Lake and Crowsnest River. Crowsnest Lake is characterized by multiple connected basins (Figure 1).^10^ The lake is fed by inflow from Crowsnest Creek, which drains a MTR waste rock pile on the Tent Mountain Mine property. A previous water quality study^8^ revealed selenium concentration as high as 24 µg/L in Crowsnest Creek downstream of the mine (at site CC1). Somewhat lower concentrations (1.3–7.0 µg/L) were noted further downstream at site CC2 where the creek empties into Crowsnest Lake.^8^ Within the lake, selenium concentrations ranged between 1.3 and 2.0 µg/L. Further downstream, Blairmore Creek and Gold Creek drain the former Grassy Mountain Mine site, and selenium concentrations in these creeks were <1 µg/L, with one exception. A ∼48-hour discharge of mine water from underground adits contained higher concentrations of selenium (2.4 µg/L) and other trace elements.^8^ These mine water discharge events have been occurring for decades, and the mine water enters the Crowsnest River. Therefore, while runoff from Tent Mountain provides a constant input of legacy selenium to the Crowsnest Lake and River system, this input is likely augmented by periods of contaminated mine water discharge into Crowsnest River from Grassy Mountain Mine.

**Figure 1.**
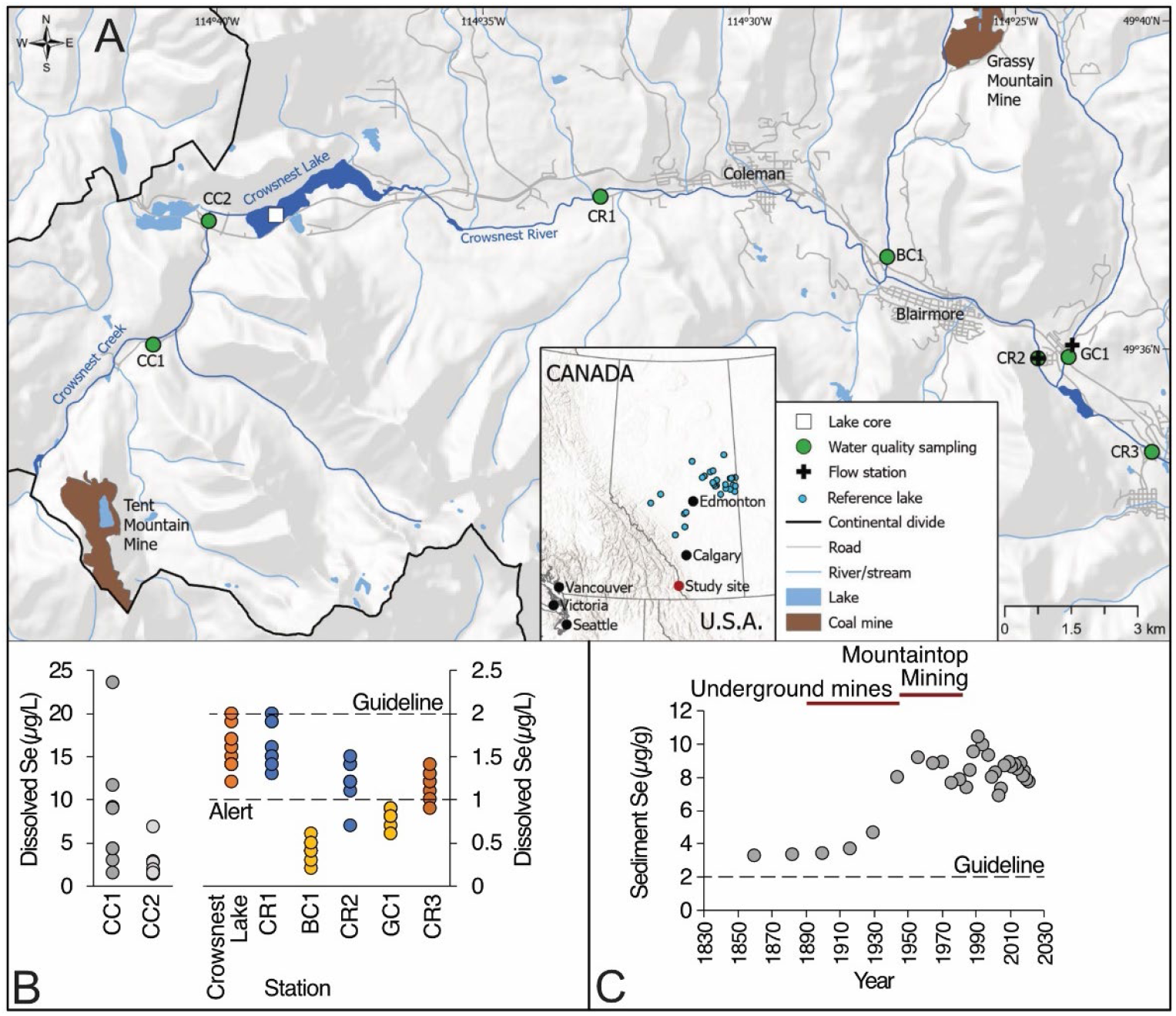
(A) Map of the Crowsnest Lake, the Crowsnest River watershed, and the various water quality sampling stations. (B) Scatter plot of selenium concentrations in Crowsnest Lake, Crowsnest River and it’s tributaries (note the different scales). Also shown are the Alert and Guideline concentrations set by the Government of Alberta.^11^ (C) Selenium concentrations (µg/g) in a sediment core from Crowsnest Lake as well as the protection of aquatic life guideline set by the Canadian Council of Ministers of the Environment (CCME). Also shown is a history of when underground and mountaintop mines were active within the Crowsnest River watershed. The water quality and sediment core data from panels B and C are from Cooke et al.^8^

### Fish tissue collection and laboratory analysis

The Crowsnest Lake fishery consists of self-sustaining populations of brown trout (*Salmo trutta*), lake trout (*Salvelinus namaycush*), and mountain whitefish (*Prosopium williamsoni*), augmented by stocking of rainbow trout (*Oncorhynchus mykiss*) and westslope cutthroat trout (*Oncorhynchus lewisi*). Historically, the lake also hosted natural populations of westslope cutthroat trout and bull trout (*Salvelinus confluentus*). Fish stocking has occurred sporadically over the past ∼100 years, and annually since 1998 (Figure S1). From 2000 to 2008, around 80,000 rainbow trout were added annually to the lake. After this date, 15,000 rainbow trout were added each year with fewer fish added in the past few years.

In August 2023, Crowsnest Lake was netted using the North American Standard Index Netting (NASIN) method for cold-water fisheries.^12^ The survey randomizes sampling sites across the lake stratified by depth. A total of 212 fish from four species were captured: brown trout (n=13), lake trout (n=21), mountain whitefish (n=71), and longnose sucker (n=107). Interestingly, no rainbow trout were caught during the survey despite the stocking history described above. Sport fish were assessed for length, weight, age, sex, and maturity (see Supporting Information for additional details), and muscle tissue samples were taken for analysis of selenium and other contaminants.

The analytical laboratories and methods used in this study align with previous assessments of metals in fish tissue in Alberta.^13–15^ Selenium and another 19 elements were measured at the Analytical and Environmental Toxicology Laboratory at the University of Alberta. In brief, ∼1 g (wet weight) of fish tissue was weighted into a microwave digestion vessel (CEM Mars 6 acid digestion system). Five milliliters of nitric acid (Optima grade) was added into each vessel, and the vessels were kept in the fume hood overnight. Five mL of deionized water was added into each vessel next day, and the vessels were loaded on the microwave digestion instrument for digestion. Each digested sample was quantitatively transferred into a 50 mL beaker and the solution evaporated to ∼1 mL at 200°C. Finally, the solution was diluted to 10 mL with 2% nitric acid for analysis. The analyses of total trace elements were conducted using an Agilent 7900 ICP-MS. Quality assurance and quality control protocols included duplicate procedural blanks during each run, re-analysis of the calibration standards every 10 to 15 samples, and comparison of measured values to certified values of standard reference material 1566b Oyster tissue (Table S1). All samples were analyzed as wet tissue (i.e., prior to any drying) and were therefore converted to dry weight using species specific moisture values (brown trout and lake trout: 78%; mountain whitefish: 80%) and standard approaches.^13,14,16^

## Results

Selenium concentrations in fish tissue samples ranged from 5 to 26 µg/g (Figure 2; Table 1). Average concentrations were highest in brown trout (18 µg/g), followed by lake trout (15 µg/g), and mountain whitefish (9 µg/g). Median selenium concentrations were significantly (Kruskal-Wallis test p<0.01) lower in mountain whitefish compared to the other two species (Table S2). Both trout species are piscivorous while the mountain whitefish feed predominately on invertebrates, which may account for the higher concentrations in the trout species.^17^ In general, fish length and weight were not strong predictors of fish tissue selenium concentrations (Figure S2).

**Table 1.** Average concentrations in µg/g dry weight and one standard deviation for elements above the method detection limit in Crowsnest Lake fish.

| Species | Se | V | Cr | Mn | Ni | Zn | As | Tl |
| --- | --- | --- | --- | --- | --- | --- | --- | --- |
| Brown trout | 18 ±4 | 0.017 ±0.005 | 0.2 ±0.1 | 0.4 ±0.1 | 0.06 ±0.02 | 24 ±6 | 0.6 ±0.8 | 0.1 ±0.04 |
| Lake trout | 15 ±2 | 0.013 ±0.005 | 0.3 ±0.5 | 0.7 ±0.2 | 0.1 ±0.05 | 20 ±2 | 0.3 ±0.4 | 0.18 ±0.09 |
| Mountain whitefish | 9 ±4 | 0.014 ±0.005 | 0.5 ±1 | 0.6 ±0.1 | 0.09 ±0.09 | 29 ±11 | 1.2 ±0.4 | 0.12 ±0.03 |

**Figure 2.**
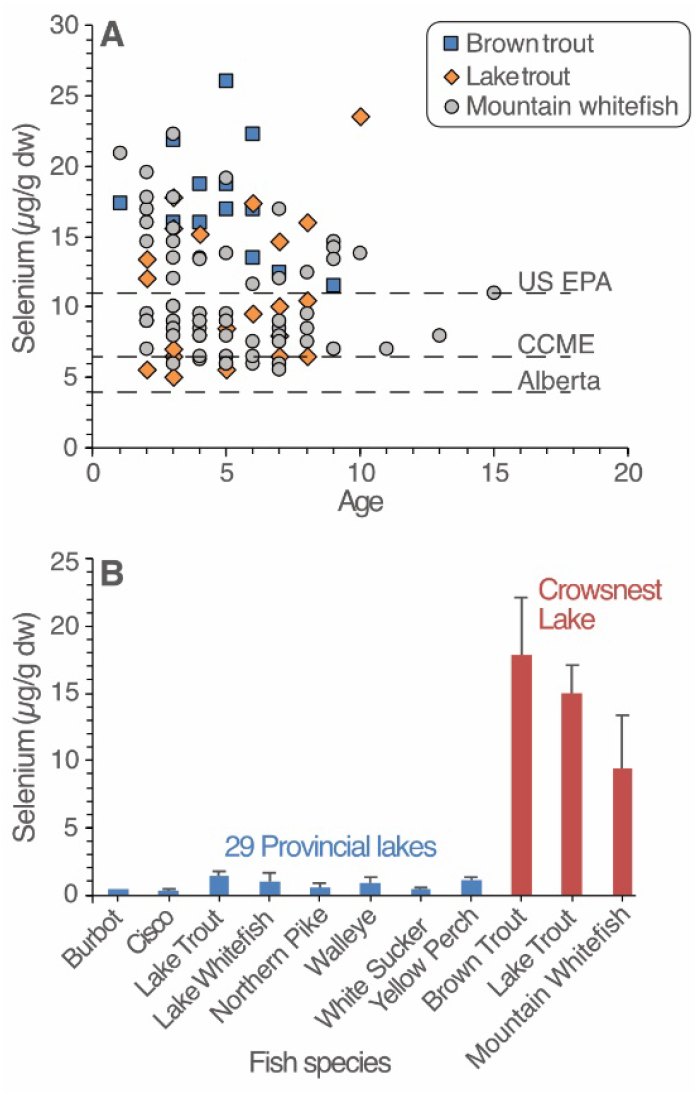
(A) Crowsnest Lake fish tissue selenium concentrations (in dry weight) and by species and fish age. Guideline values shown are for the protection of fish and wildlife consumers from direct toxicity as set by the Governments of Alberta, Canada, and the United States Environmental Protection Agency (US EPA) are also shown. (B) Average fish tissue selenium concentrations from Crowsnest Lake by species compared with species from 29 reference (i.e., not impacted by MTR coal mining) lakes from across Alberta. The error bars represent one standard deviation (SD).

Of the other elements measured, vanadium, chromium, manganese, nickel, zinc, arsenic, and thallium were above method detection limits (MDL) in all fish tissue samples (Table 1). All elemental concentration results for all samples are provided in the Supplemental Data File. Molybdenum and barium were detected in 90 and 50% of samples, respectively. Cobalt was detected in 10% of samples, while beryllium, copper, cadmium, antimony, lead, uranium, and thorium concentrations were below their respective MDL in all samples.

Alberta has an interim fish tissue selenium guideline of 4 µg/g to protect fish populations from reproductive effects.^11^ This guideline value is the same as that adopted by British Columbia’s Ministry of the Environment^18^ but lower than the Canadian Council of the Ministers of the Environment (CCME) guideline of 6.7 µg/g. Comparing against the Alberta guideline reveals 100% exceedance, meaning that every single fish sample analyzed exceeded this value (Figure 2). If we use the higher CCME guideline, we observe 84% exceedance; however, all but one of the piscivorous brown and lake trout fishes sampled exceeded the CCME and higher United States Environmental Protection Agency (US EPA) guideline of 11 µg/g.

The high fish tissue selenium concentrations observed in Crowsnest Lake are similar to those in fish populations in systems impacted by active MTR coal mining operations. For example, Kuchapski et al. reported selenium concentrations in four species of fish downstream of MTR mining operations in both the Elk River valley (in British Columbia) and the McLeod River basin (in Alberta).^17^ Across these two basins, fish populations in mining-impacted streams had an average selenium concentration of 7.6±1.0 µg/g (range: 4.7–14.3 µg/g). The Kuchapski et al. data included fish from the Fording River collected in 2011, where, less than a decade later (in 2020), the adult westslope cutthroat trout population suffered a 93% decline (relative to 2017), and juvenile density was 73% lower. The decline ultimately led to the mine owner and operator, Teck Resources Limited, being fined 60 million dollars under Canada’s *Fisheries Act* for discharges of selenium and calcite, the largest such fine in Canadian history.

Crowsnest Lake fish tissue selenium concentrations stand in contrast to fish populations from lakes not directly impacted by MTR coal mine inputs. Selenium has been measured in eight species of fish from 29 lakes across Alberta (n=840 individuals) over the past decade (Figure 2B). Within this dataset, fish tissue selenium concentrations ranged from 0.19–2.45 µg/g, well below the values observed in Crowsnest Lake fish. The contrast between Crowsnest Lake and the other Alberta lakes not impacted by MTR coal mine inputs becomes even more apparent after standardizing for fish size (Figure S3). Comparing to the provincial fish tissue dataset underscores just how anomalous the Crowsnest Lake results are.

## Discussion

The selenium concentrations and guideline exceedances noted above can only be explained by the incorporation of legacy coal mine pollution. Lake water selenium concentrations are not measured with any regularity, but sporadic measurements during the spring and summer of 2021 are presented in Figure 1B. All of the water quality results fall between 1 and 2 µg/L. These values are the Alert Concentration and a Protection of Aquatic Life Guideline, respectively, set by the Government of Alberta.^11^ Repeated measures above the Alert Concentration are intended to serve as a trigger for added research. In addition, a sediment core from Crowsnest Lake testifies that selenium supply to the lake used to be much higher (Figure 1). The core reveals that sediment selenium concentrations are now ∼4x higher than prior to the onset of regional mining activities. Even prior to mining, selenium concentrations in Crowsnest Lake bed sediment exceeded the 2 µg/g Protection of Aquatic Life Guideline set by the CCME.^19^

The source of this selenium seems to be periodic inputs of high-selenium water from Crowsnest Creek, which drains the legacy Tent Mountain Mine. Runoff from the mine waste-rock drains into Crowsnest Creek, and a 2021 assessment revealed selenium concentrations up to 24 µg/L in the creek (Figure 1B). Crowsnest Creek flows into Crowsnest Lake, where a lake water residence time of ∼6 months^10^ would offer opportunity for selenium uptake by biota. Previous research has revealed that fish occupying lakes (or even oxbow lakes) often contain higher selenium concentrations than fish residing in rivers or streams. This is because slow-water lentic systems offer prime feeding locations and serve as a deep-water refuge during drought or seasonal periods of low flow. These characteristics result in higher bioaccumulation and associated toxic hazard. This appears to be the same mechanism at play in Crowsnest Lake. Measuring selenium in foodweb organisms at various trophic levels could reveal insight into the factors driving biomagnification in this system.

We are only aware of one previous assessment of fish tissue selenium within the Crowsnest River system. In 2016, eight fish from both Blairmore and Gold Creeks were analyzed for tissue selenium as part of the Grassy Mountain Coal Project Environmental Impact Assessment.^20^ The species analyzed included brook trout (*Salvelinus fontinalis*), rainbow trout, and a hybrid population of rainbow trout and westslope cutthroat trout. Selenium concentrations in these individuals ranged from 4.5 to 10.4 µg/g (Table S3). While these fish tissue concentrations are not as high as those in Crowsnest Lake, they suggest a broader selenium problem within the Crowsnest Lake and River watershed.^1^

There is no flow control on the outlet of Crowsnest Lake (or on any of the tributaries flowing into the lake), which allows fish migration within the Crowsnest River system. Selenium concentrations in the Crowsnest Lake fishery are high enough that we might expect clinical expression of selenium poisoning, which can include behavioral changes, physical symptoms (e.g., edemas and fin and tail damage), respiratory issues, reproductive issues, and ultimately population collapse. Clinical expression of selenium poisoning was not observed during our assessment; however, fish with some of these symptoms have been observed previously in the Crowsnest River. James et al. conducted a 2019 assessment of the Crowsnest River in response to rainbow trout population declines due to Whirling Disease.^21^ The authors noted that 85% of captured wild fingerling rainbow trout exhibited clinical signs of disease. Whirling Disease and excess selenium poisoning share many of the same clinical symptoms, including deformities, lethargy, and potential mortality especially in early life stages. Thus, it remains possible that some of the clinical symptoms observed by James et al. could have been from selenium poisoning.

Crowsnest Lake is a managed recreational fishery. Over one million rainbow trout have been added to Crowsnest Lake since 1998 (Figure S2), yet we did not net a single rainbow trout in 2023. This is even though fish introduced between 1998 and 2009 (or 83% of all fish added since 1998) were capable of reproduction (i.e., 2N strain), and fish were stocked into the lake three months prior to netting. Additional research is therefore required to understand fish population dynamics in this system. Moreover, while rainbow trout in the river have been assessed for Whirling Disease^21^, no such assessment has been made for selenium – our results emphasize this should be done.

In addition to the toxicity risks to fish, there are also concerns for human health for those that consume fish high in selenium. In the United States, states with active MTR coal mines have issued fish consumption advisories to limit the intake of fish high in coal mine selenium pollution. This includes the Mudd River in West Virginia, where decades of research have documented elevated fish tissue selenium concentrations downstream of active MTR mines.^22^ Current (2024) fishing regulations in Crowsnest Lake allow for the harvesting of any size of brown and lake trout but only mountain whitefish longer than 300 mm can be kept by anglers.^23^ Given the sportfish we measured had selenium concentrations comparable to the fisheries mentioned above in the United States, consumption advisories for the Crowsnest Lake and Crowsnest River system seem warranted.

Crowsnest Lake and its outlet, the Crowsnest River, are highly stressed aquatic ecosystems. In addition to the emergence of Whirling Disease^21^, declining snowpack and increased drought,^24^ year-round recreational angling pressure^23^, and now our new evidence for elevated selenium in fish, make the Crowsnest Lake and River an especially vulnerable system. Any new development of coal mining along the eastern slopes may well push the Crowsnest fishery beyond recovery.

## Supporting Information

Summary of fish netting methods and result; graph of fish stocked in Crowsnest Lake; plots of selenium concentrations in fish tissue versus fish metrics; fish tissue to length ratios; tables of reference material recoveries, Kruskal-Wallis p-values, and additional elemental concentrations.

## Acknowledgements

The authors thank field staff from the Government of Alberta for conducting the survey and collecting the fish used in this study. Funding for this work was provided by the Government of Alberta.

## Summary of fish netting methods and results

The American Fisheries Society proposed gill nets (North American Standard Index Netting (NASIN)) as the standard for community-based sampling of angler harvested freshwater species in North America.^1^ Index netting was conducted from August 14–17, 2023. Twelve “core-mesh” gill nets (i.e., NASIN nets) were set at random locations between 1.8 and 20 meters deep for 17–19 hours (i.e., a net-night), and then reset in new random locations. The information collected from sport fish includes fish length, weight, age, sex, and maturity.

### Brown Trout

The mean catch rate of brown trout was 1.08/fish/net-night (95% Confidence Interval (CI): 0–0.17). The brown trout in Crowsnest Lake display a normal size and age-class distributions with the female brown trout (n=6) ranging in age from 2 to 6 years-old, reaching sexual maturity at approximately age 5 and ranged in size from 263–505 mm total length. Male brown trout (n=7) ranged in age from 1 to 7 years-old, reaching sexual maturity at approximately age 3 and ranged in size from 216–538 mm total length.

### Lake Trout

The mean catch rate of lake trout was 1.75 fish/net-night (95% CI: 0.71–2.78). The lake trout in Crowsnest Lake display a wide size and age-class distributions with the female lake trout (n=7) ranging in age from 1 to 15 years-old, unknown age at which they approximately reach sexual maturity because of small sample size (n=1) of mature fish, but we collected fish ranging between 205–760 mm total length. Male lake trout (n=6) ranged in age from 2 to 7 years-old, but we were not able to confirm that we collected a single sexual maturity male and ranged in size from 260–533 mm total length.

### Mountain Whitefish

The mean catch rate of mountain whitefish was 5.91 fish/net-night (95% CI: 3.16–8.65). The mountain whitefish in Crowsnest Lake display a wide size and age-class distributions with the female mountain whitefish (n=37) ranging in age from 3 to 13 years-old reaching sexual maturity at approximately age 7 and ranged in size from 188–357 mm total length. Male mountain whitefish (n=30) ranged in age from 2 to 9 years-old reaching sexual maturity at approximately age 5 and ranged in size from 174–275 mm total length.

### Summary

Mountain whitefish from 174–357 mm total length made up the largest part of the total catch in Crowsnest Lake in 2023. Brown trout and lake trout were also present, but in smaller numbers than observed for mountain whitefish. Additionally, we collected many longnose sucker. However, despite extensive annual stocking totalling more than 210,000 black spotted trout dating back to 2009, (both hatchery derived rainbow trout and westslope cutthroat trout) not a single rainbow trout or westslope cutthroat trout was caught during this survey.

Based on the variety of size classes, there appears to be limited but consistent lake trout recruitment occurring in Crowsnest Lake, with the majority of captured fish immature. Mountain whitefish and brown trout continue to utilize the lake to a degree, but it is possible they are also utilizing the Crowsnest River for portions of the year.

---

**Figure S1.**
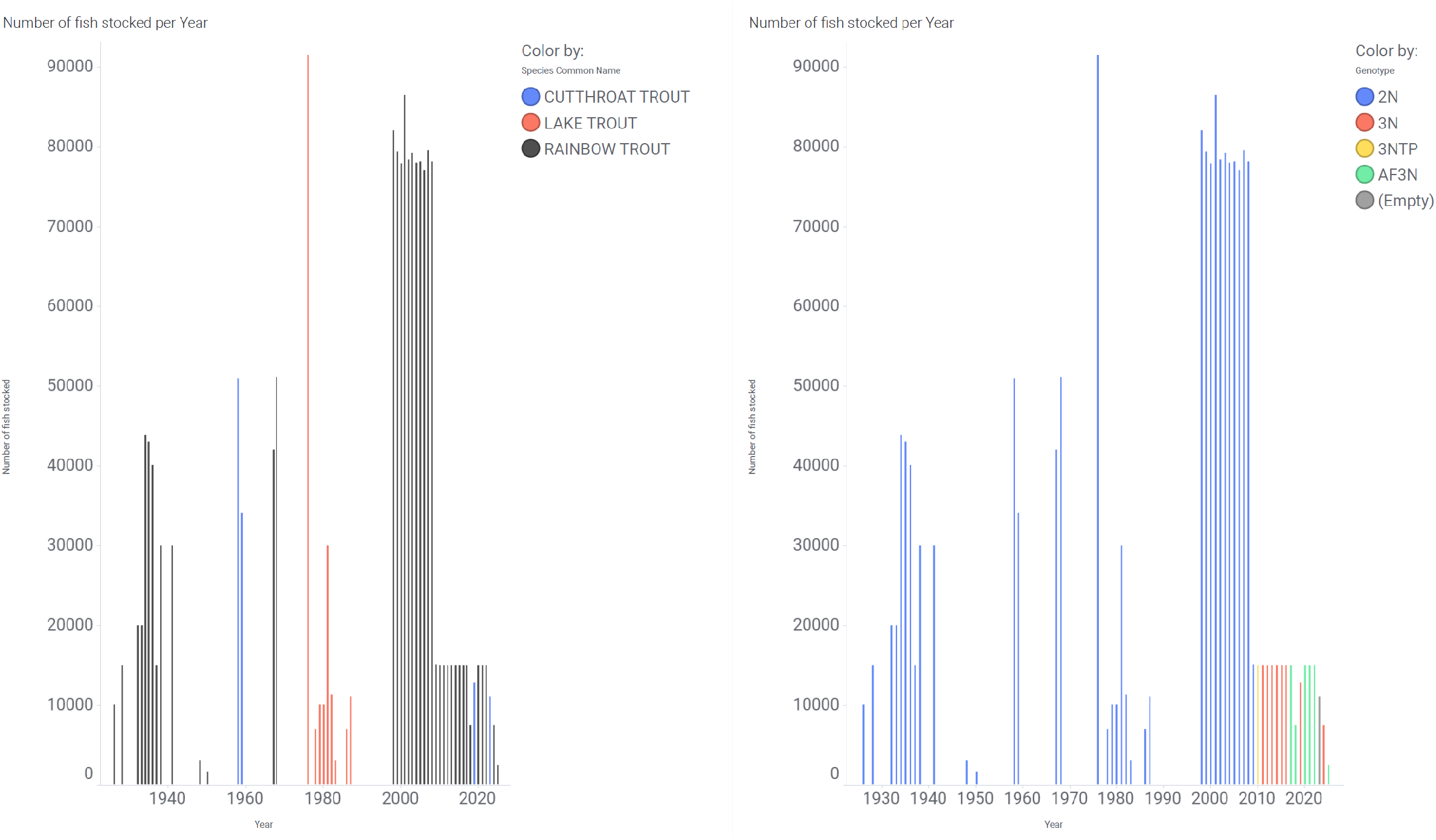
Number of fish stocked in Crowsnest Lake by species (left) and by genotype (right) for the past ∼100 years.

**Figure S2.**
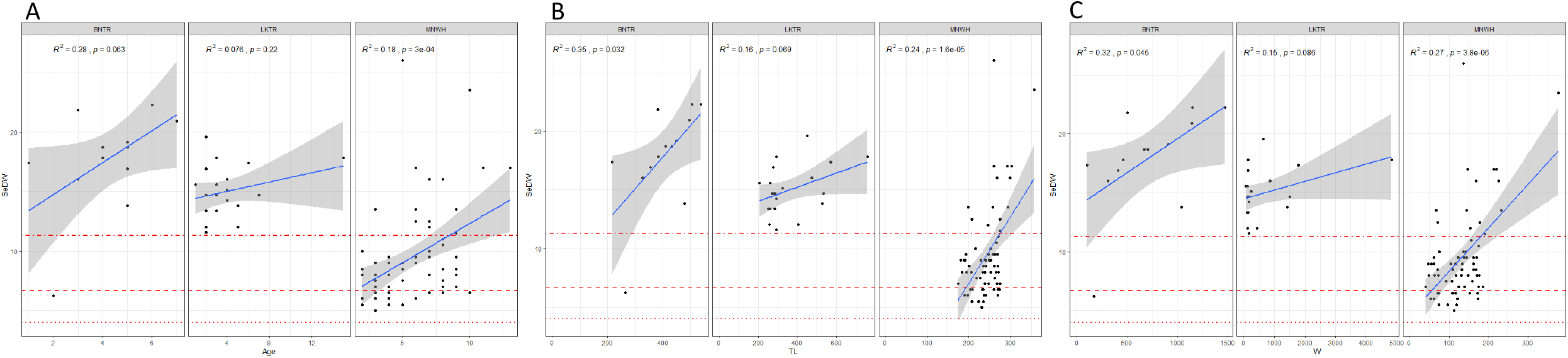
Plots of selenium concentrations in dry weight (SeDW) versus (A) fish age, (B) fish total length (TL) in mm, and (C) fish weight (W) in grams.

**Figure S3.**
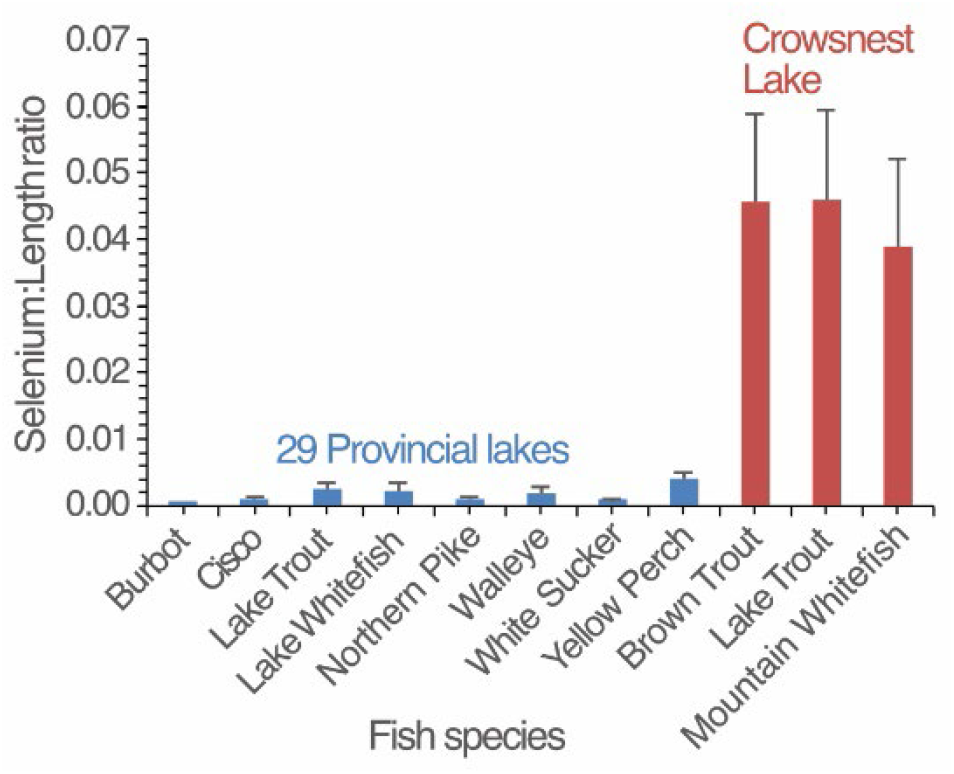
Average and 1 SD selenium to length ratio in fish from 29 provincial lakes compared with results from Crowsnest Lake.

**Table S1.**
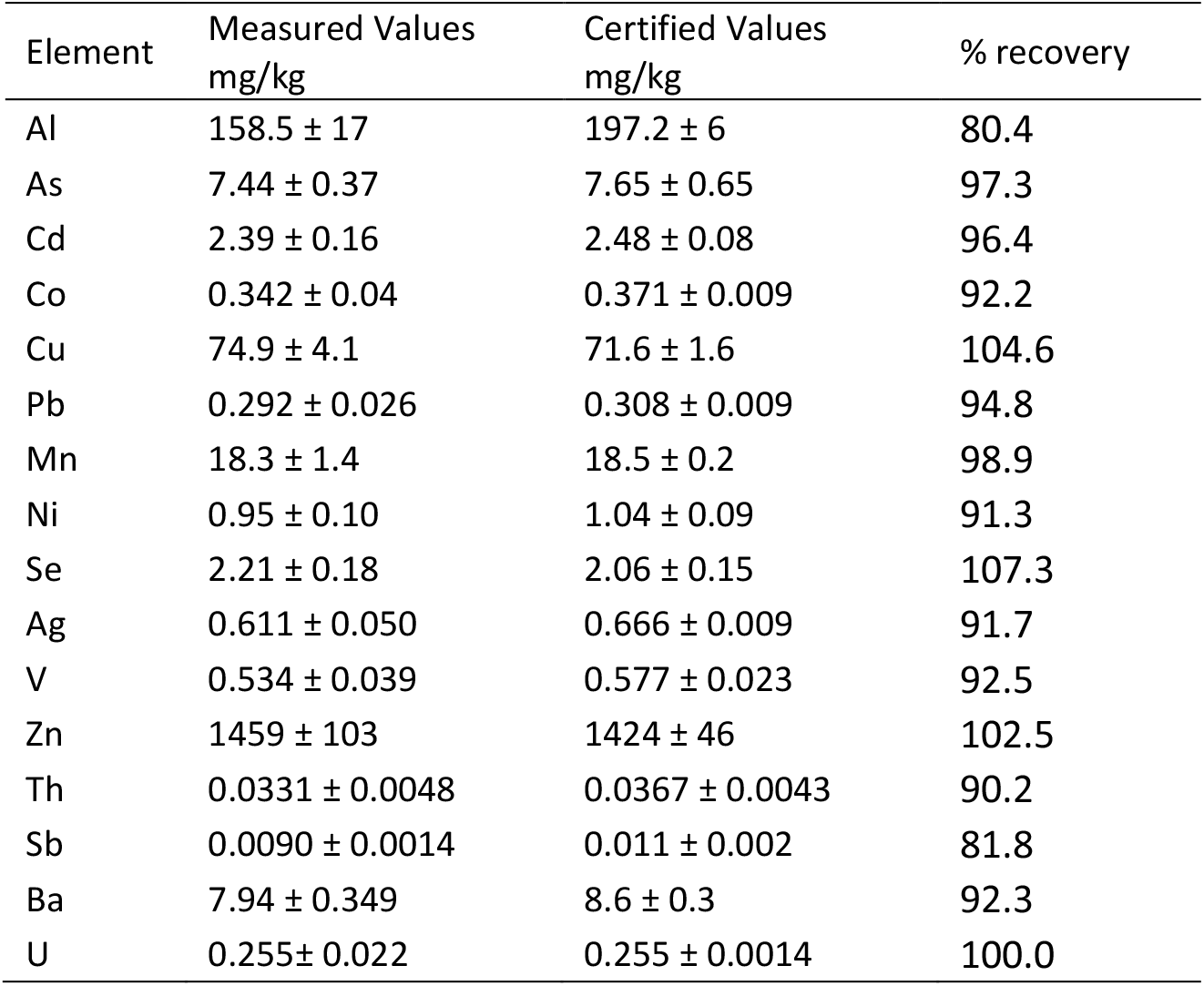
Concentrations of elements in SRM standard reference material 1566b Oyster tissue and the certified values.

**Table S2.** P-values from a Kruskal-Wallis test for equal medians.

| Species | Brown trout | Lake trout | Mountain whitefish |
| --- | --- | --- | --- |
| Brown trout |  | 0.40 | 4.03E-08 |
| Lake trout | 0.40 |  | 4.69E-08 |
| Mountain whitefish | 4.03E-08 | 4.69E-08 |  |

**Table S3.** Average and one standard deviation elemental concentrations in µg/g wet weight in brown trout, lake trout, and mountain whitefish from Crowsnest Lake.

| Species | Se | V | Cr | Mn | Ni | Zn | As | Tl |
| --- | --- | --- | --- | --- | --- | --- | --- | --- |
| Brown trout | 4 ±1 | 0.004 ±0.001 | 0.04 ±0.03 | 0.09 ±0.02 | 0.014 ±0.003 | 5.4 ±1.3 | 0.13 ±0.17 | 0.02 ±0.01 |
| Lake trout | 3.4 ±0.5 | 0.003 ±0.001 | 0.07 ±0.12 | 0.15 ±0.04 | 0.023 ±0.012 | 4.4 ±0.5 | 0.06 ±0.09 | 0.04 ±0.02 |
| Mountain whitefish | 1.9 ±0.8 | 0.003 ±0.001 | 0.11 ±0.2 | 0.12 ±0.03 | 0.018 ±0.019 | 5.8 ±2.2 | 0.24 ±0.09 | 0.02 ±0.01 |

## References

(1) Palmer, M. A.; Bernhardt, E. S.; Schlesinger, W. H.; Eshleman, K. N.; Foufoula-Georgiou, E.; Hendryx, M. S.; Lemly, A. D.; Likens, G. E.; Loucks, O. L.; Power, M. E.; White, P. S.; Wilcock, P. R. Mountaintop Mining Consequences. Science. 2010, 327 (5962), 148–149. 10.1126/science.1180543.

(2) Nippgen, F.; Ross, M. R. V.; Bernhardt, E. S.; McGlynn, B. L. Creating a More Perennial Problem? Mountaintop Removal Coal Mining Enhances and Sustains Saline Baseflows of Appalachian Watersheds. Environ. Sci. Technol. 2017, 51 (15), 8324–8334. 10.1021/acs.est.7b02288.

(3) Cooke, C. A.; Drevnick, P. E. Transboundary Atmospheric Pollution from Mountaintop Coal Mining. Environ. Sci. Technol. Lett. 2022, 9 (11), 943–948. 10.1021/acs.estlett.2c00677.

(4) Palace, V. P.; Baron, C.; Evans, R. E.; Holm, J.; Kollar, S.; Wautier, K.; Werner, J.; Siwik, P.; Sterling, G.; Johnson, C. F. An Assessment of the Potential for Selenium to Impair Reproduction in Bull Trout, Salvelinus Confluentus, from an Area of Active Coal Mining. Environ. Biol. Fishes 2004, 70 (2), 169–174. 10.1023/B:EBFI.0000029346.60306.34.

(5) Wayland, M.; Crosley, R. Selenium and Other Trace Elements in Aquatic Insects in Coal Mine-Affected Streams in the Rocky Mountains of Alberta, Canada. Arch. Environ. Contam. Toxicol. 2006, 50 (4), 511–522. 10.1007/s00244-005-0114-8.

(6) Lindberg, T. T.; Bernhardt, E. S.; Bier, R.; Helton, A. M.; Merola, R. B.; Vengosh, A.; Di Giulio, R. T. Cumulative Impacts of Mountaintop Mining on an Appalachian Watershed. Proc. Natl. Acad. Sci. 2011, 108 (52), 20929–20934. 10.1073/pnas.1112381108.

(7) Storb, M. B.; Bussell, A. M.; Caldwell Eldridge, S. L.; Hirsch, R. M.; Schmidt, T. S. Growth of Coal Mining Operations in the Elk River Valley (Canada) Linked to Increasing Solute Transport of Se, NO 3 –, and SO 4 2– into the Transboundary Koocanusa Reservoir (USA–Canada). Environ. Sci. Technol. 2023, 57 (45), 17465–17480. 10.1021/acs.est.3c05090.

(8) Cooke, C. A.; Emmerton, C. A.; Drevnick, P. E. Legacy Coal Mining Impacts Downstream Ecosystems for Decades in the Canadian Rockies. Environ. Pollut. 2024, 344, 123328. 10.1016/j.envpol.2024.123328.

(9) Brooks, A. C.; Ross, M. R. V.; Nippgen, F.; McGlynn, B. L.; Bernhardt, E. S. Excess Nitrate Export in Mountaintop Removal Coal Mining Watersheds. J. Geophys. Res. Biogeosciences 2019, 124 (12), 3867–3880. 10.1029/2019JG005174.

(10) Mitchell, P.; Prepas, E. Atlas of Alberta Lakes; The University of Alberta Press: Edmonton, 1990.

(11) Goverment of Alberta. Environmental Quality Guidelines for Alberta Surface Waters; Edmonton, Alberta, 2018. https://open.alberta.ca/publications/9781460138731.

(12) Standard Methods for Sampling North American Freshwater Fishes; Bonar, S. A., Hubert, W. A., Willis, D. W., Eds.; American Fisheries Society, 2009. 10.47886/9781934874103.fmatter.

(13) Donadt, C.; Cooke, C. A.; Graydon, J. A.; Poesch, M. S. Biological Factors Moderate Trace Element Accumulation in Fish along an Environmental Concentration Gradient. Environ. Toxicol. Chem. 2021, 40 (2). 10.1002/etc.4926.

(14) Donadt, C.; Cooke, C. A.; Graydon, J. A.; Poesch, M. S. Mercury Bioaccumulation in Stream Fish from an Agriculturally-Dominated Watershed. Chemosphere 2021, 262, 128059. 10.1016/j.chemosphere.2020.128059.

(15) Emmerton, C. A.; Drevnick, P. E.; Serbu, J. A.; Cooke, C. A.; Graydon, J. A.; Reichert, M.; Evans, M. S.; McMaster, M. E. Downstream Modification of Mercury in Diverse River Systems Underscores the Role of Local Conditions in Fish Bioaccumulation. Ecosystems 2023, 26 (1), 114–133. 10.1007/s10021-022-00745-w.

(16) Scudder, B. C.; Chasar, L. C.; Wentz, D. A.; Bauch, N. J.; Brigham, M. E.; Moran, P. W.; Krabbenhoft, D. P. Mercury in Fish, Bed Sediment, and Water from Streams across the United States, 1998-2005. In Contaminated Fish: Chemical Residues and Mercury; 2011.

(17) Kuchapski, K. A.; Rasmussen, J. B. Food Chain Transfer and Exposure Effects of Selenium in Salmonid Fish Communities in Two Watersheds in the Canadian Rocky Mountains. Can. J. Fish. Aquat. Sci. 2015, 72 (7). 10.1139/cjfas-2014-0484.

(18) Government of British Columbia. Ambient Water Quality Guidelines for Selenium; 2014.

(19) CCME. Canadian Sediment Quality Guidelines for the Protection of Aquatic Life; Winnipeg, MB, 2001.

(20) Benga Mining Limited. Grassy Mountain Coal Project Environmental Impact Assessment Section B: Geology and Geotechnical; Ottawa, Canada, 2016. https://iaac-aeic.gc.ca/050/documents/p80101/115589E.pdf.

(21) James, C. T.; Veillard, M. F.; Martens, A. M.; Pila, E. A.; Turnbull, A.; Hanington, P.; Luek, A.; Alexander, J.; Nehring, R. B. Whirling Disease in the Crowsnest River: An Emerging Threat to Wild Salmonids in Alberta. Can. J. Fish. Aquat. Sci. 2021, 78 (12), 1855–1868. 10.1139/cjfas-2020-0484.

(22) West Virginia DNR. Fishing Regulations Summary. https://dhhr.wv.gov/News/2024/Pages/2024-Sport-Fishing---New-Change-to-the-2023-West-Virginia-Fish-Consumption-Advisories.aspx (accessed 2025-02-13).

(23) Government of Alberta. 2024 Alberta Guide to Sportfishing Regulations; Edmonton, 2024. https://open.alberta.ca/publications/alberta-guide-to-sportfishing-regulations.

(24) Newton, B. W.; Farjad, B.; Orwin, J. F. Spatial and Temporal Shifts in Historic and Future Temperature and Precipitation Patterns Related to Snow Accumulation and Melt Regimes in Alberta, Canada. Water 2021, 13 (8), 1013. 10.3390/w13081013.

## References cited in the Supporting Information

(1) Standard Methods for Sampling North American Freshwater Fishes; Bonar, S. A., Hubert, W. A., Willis, D. W., Eds.; American Fisheries Society, 2009. 10.47886/9781934874103.fmatter.

(2) Donadt, C.; Cooke, C. A.; Graydon, J. A.; Poesch, M. S. Biological Factors Moderate Trace Element Accumulation in Fish along an Environmental Concentration Gradient. Environ. Toxicol. Chem. 2021, 40 (2). 10.1002/etc.4926.

